# Detection and discrimination of influenza B Victoria lineage deletion variant viruses by real-time RT-PCR

**DOI:** 10.1101/818617

**Authors:** Bo Shu, Marie K. Kirby, Christine Warnes, Wendy M. Sessions, William G. Davis, Ji Liu, Malania M. Wilson, David E. Wentworth, John R. Barnes

**Affiliations:** Virology, Surveillance and Diagnostic Branch, Influenza Division, Centers for Disease Control and Prevention, Atlanta, USA; Battelle Memorial Institute, Atlanta, USA; Chickasaw Nation Industries, Inc., Atlanta, USA

## Abstract

Influenza B viruses have two genetically and antigenically distinct lineages, B/Victoria/2/1987-like (VIC) and B/Yamagata/16/1988-like (YAM) viruses, that emerged in the 1980s and co-circulate annually during the influenza season. During the 2016-2017 influenza season, influenza B/VIC lineage variant viruses emerged with two (K_162_N_163_) or three (K_162_N_163_D_164_) amino acid (AA) deletions in the hemagglutinin protein. Hemagglutination inhibition assays demonstrate that these deletion variant influenza B/VIC viruses are antigenically distinct from each other and from the progenitor B/VIC virus that lacks the deletion. Therefore, there are currently four antigenically distinct HA proteins expressed by influenza B co-circulating: B/YAM, B/VIC V1A (no deletion), B/VIC V1A.1 (two-AA deletion), and B/VIC V1A.2 and V1A.3 (three-AA deletion). The prevalence of these viruses differs across geographic regions, making it critical to have a sensitive, rapid diagnostic assay(s) that detect and distinguish these Influenza B variant viruses during surveillance. Here, we present a real time RT-PCR assay that targets the influenza B/VIC deletion region in the HA gene and detects and distinguishes the influenza B/VIC V1A, B/VIC V1A.1, B/VIC V1A.2 and B/VIC V1A.3 variant viruses, with no cross-reactivity. This assay can be run as a multiplex reaction, allowing for increased testing efficiency and reduced cost. Coupling this assay with the CDC Human Influenza Virus Real-Time RT-PCR Diagnostic Panel Influenza B Lineage Genotyping Kit results in rapid detection and characterization of circulating influenza B viruses. Having accurate and detailed surveillance information on these distinct Influenza B variant viruses will provide insight into the prevalence and geographic distribution and could aid in vaccine recommendations.

## INTRODUCTION

Influenza B viruses co-circulate with influenza A strains during the annual influenza season, contributing to overall mortality and morbidity of influenza epidemics. In the 1980’s, influenza B viruses evolved into two genetically and antigenically distinct lineages represented by B/Victoria/2/1987 (VIC) and B/Yamagata/16/1988 (YAM), that co-circulate in human beings during influenza seasons worldwide [1–5]. During the 2016-2017 influenza season, the Centers for Disease Control and Prevention (CDC) detected influenza B/VIC viruses in the United States (U.S) that were antigenically distinct from the World Health Organization (WHO) recommended vaccine virus, B/Brisbane/60/2008 (VIC)[6]. Genetic analysis confirmed these viruses have a deletion of six nucleotides in the hemagglutinin (HA) gene resulting in a two amino acid (AA) deletion at positions 162 and 163 (corresponding to nucleotide positions 529-534)[6]. This B Victoria lineage two AA deletion virus (V1A.1; K_162_N_163_ deletion) has since spread and been detected worldwide. Two B/Vic lineage variants with three AA deletions (V1A.2 and V1A.3) at AA positions 162-164 (K_162_N_163_D_164_, and nucleotide positions 529-537) emerged shortly after and subsequently via parallel evolution. The 3DEL viruses have since been detected in Asia, Africa, Europe and America [7]. Hemagglutination inhibition (HI) assays demonstrated that both the V1A.1 and V1A 3DEL variant influenza B viruses are antigenically distinct from each other and to the no-deletion B/VIC virus (V1A) (Table 1). Thus, there are currently five genetically distinct HA genes that yield four antigenically distinct Influenza B viruses currently co-circulating: B/YAM, B/VIC V1A, B/VIC V1A.1, B/VIC V1A.2 and B/VIC V1A.3 (Table 1; Figure 1).

**Table 1.** Hemagglutination inhibition reactions of Influenza B viruses

| Influenza B viruses strain | Ferret antiserum |  |  |  |  |
| --- | --- | --- | --- | --- | --- |
|  | B/Brisbane/60/2008<br>(V1A) | B/Maryland/15/2016<br>(V1A.1) | B/Hong Kong/286/2017<br>(V1A.2) | B/Washington/02/2019<br>(V1A.3) | B/Phuket/3073/2013<br>(Yam) |
| V1A |  |  |  |  |  |
| **B/Brisbane/60/2008** | **2560** | 10 | 40 | 40 | < 10 |
| V1A.1 |  |  |  |  |  |
| **B/Maryland/15/2016** | 80 | **160** | 10 | 20 | < 10 |
| **B/Colorado/06/2017** | 80 | 160 | 10 | 10 | < 10 |
| V1A.2 |  |  |  |  |  |
| B/Hong Kong/286/2017 | 320 | 40 | **320** | **80** | < 10 |
| **B/Hong Kong/269/2017** | 80 | 20 | 160 | 40 | < 10 |
| V1A.3 |  |  |  |  |  |
| **B/Washington/02/2019** | 40 | 40 | 40 | **320** | < 10 |
| Yam |  |  |  |  |  |
| **B/Phuket/3073/2013** | < 10 | < 10 | < 10 | < 10 | **640** |

Viruses used in this study are shown in bold and numbers in bold indicate homologous HI titers

**Figure 1.**
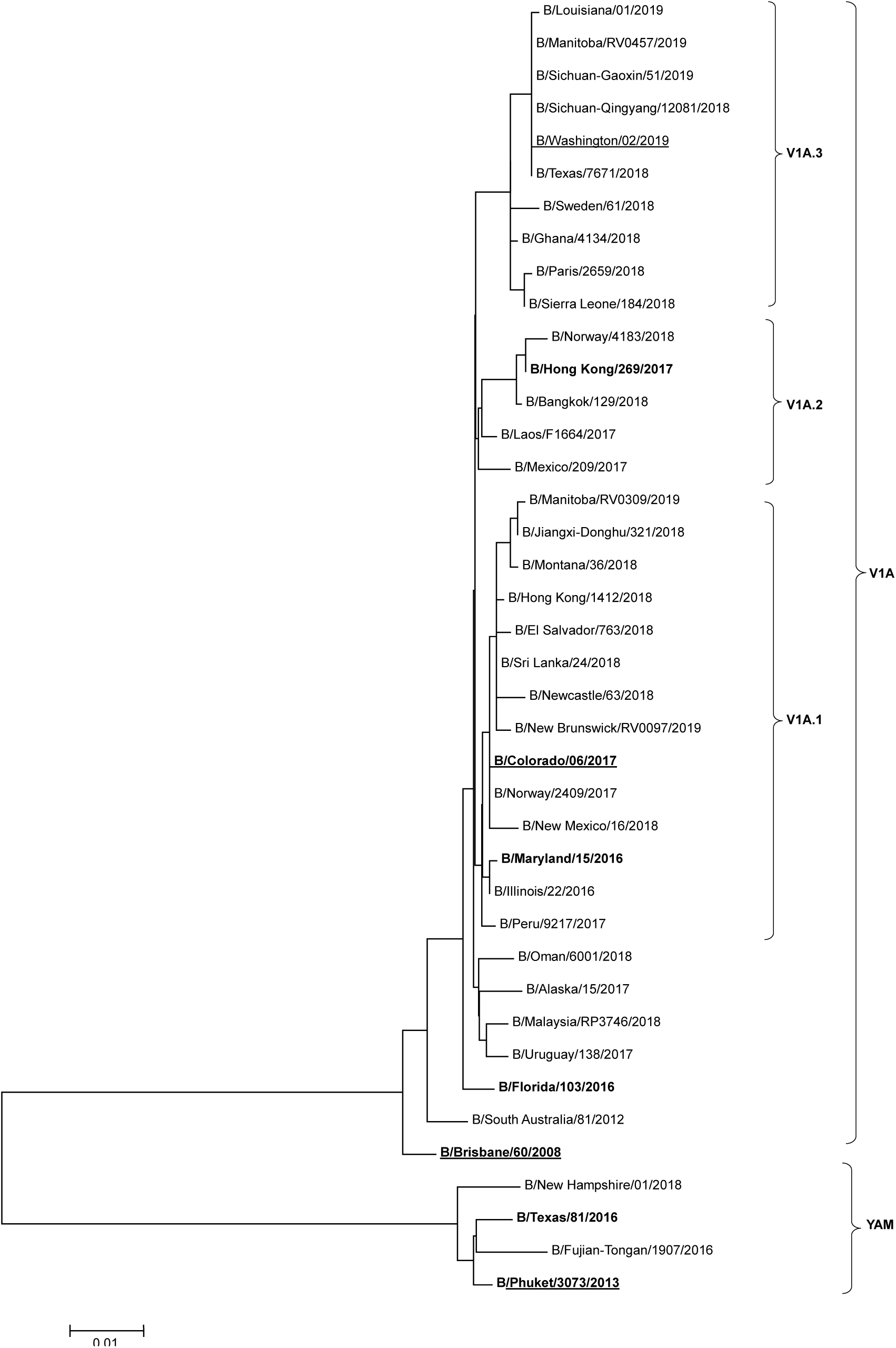
Evolutionary relationships among Influenza B Victoria and Yamagata Hemagglutinin genes. HA phylogenetic tree with representative viruses of the four genetic groups of B/Victoria and B/Yamagata lineages of related viruses; V1A, V1A.1, V1A.2, V1A.3 and YAM indicated by the bars on the right. The viruses evaluated in this study are showed in bold and influenza B vaccine viruses are underlined. Phylogenetic analysis was performed using Molecular Evolutionary Genetics Analysis software (MEGA, version 7.0). The evolutionary history was inferred using the Neighbor-Joining method.

The most effective method for prevention and control of influenza infection is vaccination [3, 4, 8]. Licensed seasonal vaccines are updated annually and the WHO makes recommendations on the composition of influenza virus vaccines on the basis of surveillance, laboratory, and clinical observations [3]. This process occurs twice a year, in February for the northern hemisphere and in September for the southern hemisphere. It is important to identify the most prevalent influenza A subtype and B lineage viruses through influenza surveillance for influenza vaccine selection, especially for people who may not have been exposed to a subtype/lineage of influenza virus [9]. Identifying the optimal viruses to include in the next season’s influenza vaccines is a significant challenge for the influenza surveillance network known as the Global Influenza Surveillance and Response System (GISRS). A diagnostic assay that allows for rapid identification of these genetically and antigenically distinct Influenza B viruses is beneficial to understanding how the prevalence of these viruses varies within different geographic regions of the world and will provide data that could be leveraged to guide relevant vaccine strain selection.

A real time RT-PCR (rRT-PCR)-based B/Victoria lineage deletion detection assay (Vic deletion assay) for detection and discrimination of these antigenically drifted B/Vic deletion variant viruses is detailed herein. The Vic deletion assay consists of a set of conserved primers and deletion-specific dual-labeled hydrolysis probes and has high sensitivity and specificity to distinguish between V1A, V1A.1 V1A.2 and V1A.3 groups without cross-reactivity, allowing for multiplex reactions for maximum testing efficiency. This assay is currently the only available method, to our knowledge, that can distinguish these four B/VIC genetic groups, outside of a pyrosequencing technique and a conventional RT-PCR method, both recently published [10, 11]. Real-time RT-PCR equipment is readily available in the majority of diagnostic and surveillance laboratories, it is highly sensitive, specific, and extremely rapid, making it the optimal assay for diagnostic and surveillance purposes. When used in conjunction with the CDC Human Influenza Virus Real-Time RT-PCR Diagnostic Panel (CDC Flu rRT-PCR Dx Panel) Influenza B Lineage Genotyping Kit, all five influenza B genetic backgrounds (B/Vic V1A, V1A.1, V1A.2, V1A.3 and B/YAM) can be distinguished in the circulating influenza viruses, allowing for identification of the predominating influenza B virus within specific populations and global regions, which in turn will aid in shaping vaccine recommendations for future influenza seasons.

## MATERIALS AND METHODS

### Influenza Viruses, Antigenic analysis and RNA Extraction

Influenza viruses tested in this study were grown to high titer in either Madin-Darby Canine Kidney (MDCK) cells or embryonated chicken eggs (ECE) [12]. Infectious virus in culture supernatants or allantoic fluids was measured by tissue culture infectious dose-50% (TCID_50_/ml) or egg-infectious dose-50% (EID_50_ /ml), respectively[13]. Influenza B virus isolates used for analytical performance evaluation included B/Maryland/15/2016 and B/Colorado/6/2017 (V1A.1), B/Hong Kong/269/2017(V1A.2) and B/Washington/02/2019 (V1A.3) as well as no deletion viruses B/Florida/103/2016 and B/Brisbane/60/2008 (V1A). YAM lineage viruses represented by B/Phuket/3073/2013 and B/Texas/81/2016 were also included.

Influenza B virus lineages were confirmed using antigenic characterization by HI assay on MDCK-propagated virus isolates and genetic sequence analysis (Table 1, Figure 1).

The antigenic characteristics of virus isolates were determined by HI tests using post-infection antisera. The HI test was performed as described previously (Kendal and Cate, 1983)[14].

Viral RNA was extracted from 100µl of supernatant or allantoic fluid and eluted into 100µl of RNA elution buffer using MagNA Pure Compact RNA isolation kit on a MagNA Pure Compact instrument (Roche Applied Science) [15].

### The B/Victoria lineage deletion detection assay primers/probes

Oligonucleotide primers and probes of Vic deletion assay were designed based on available nucleotide sequence data from GenBank database of National Centers for Biological Information (NCBI) and the Global Initiative on Sharing Avian Influenza Data (GISAID). The Vic deletion assay includes a single set of conserved amplification primers and three deletion-specific dual-labeled hydrolysis probes, including VIC 2_Del, Vic 3_Del and Vic No_Del probes. Three probes, targeted on deletion region of the HA gene of influenza B viruses, were designed to specifically detect and differentiate V1A.1, V1A 3DEL and the V1A genetic group (No deletion) viruses (B/Vic No_Del), respectively (Figure S1); Vic 2_Del and Vic No_Del probes were designed using BHQ*plus*^TM^ dual-labeled hydrolysis probes (BHQplus) that were labeled at the 5’-end with the reporter molecule 6-carboxyfluorescein (FAM) or Hex and CAL Fluor Red 610 and with Blackhole Quencher™ 1 (BHQ™1) (FAM or Hex) or BHQ™2 (CAL Fluor Red 610) at the 3’-end. The Vic 3_Del probe was labeled at the 5’-end with the reporter molecule 6-carboxyfluorescein (FAM) and (BHQ™) 1 at the 3’-end, and included a triplet of Locked Nucleic Acids (LNA) [16, 17] that centered on the mismatch bases from V1A.1 and V1A viruses.

Primer probe sequence specificity was also evaluated by sequence analysis of 51,759 gene segments available in NCBI or GISAID databases. Primers were designed to have annealing temperatures of approximately 60°C and probes were designed to have annealing temperatures of approximately 68°C using PrimerExpress 3.0 software (Applied Biosystems, Foster City, USA). Primers were synthesized by the Biotechnology Core Facility at the CDC (Table 2).

**Table 2:**
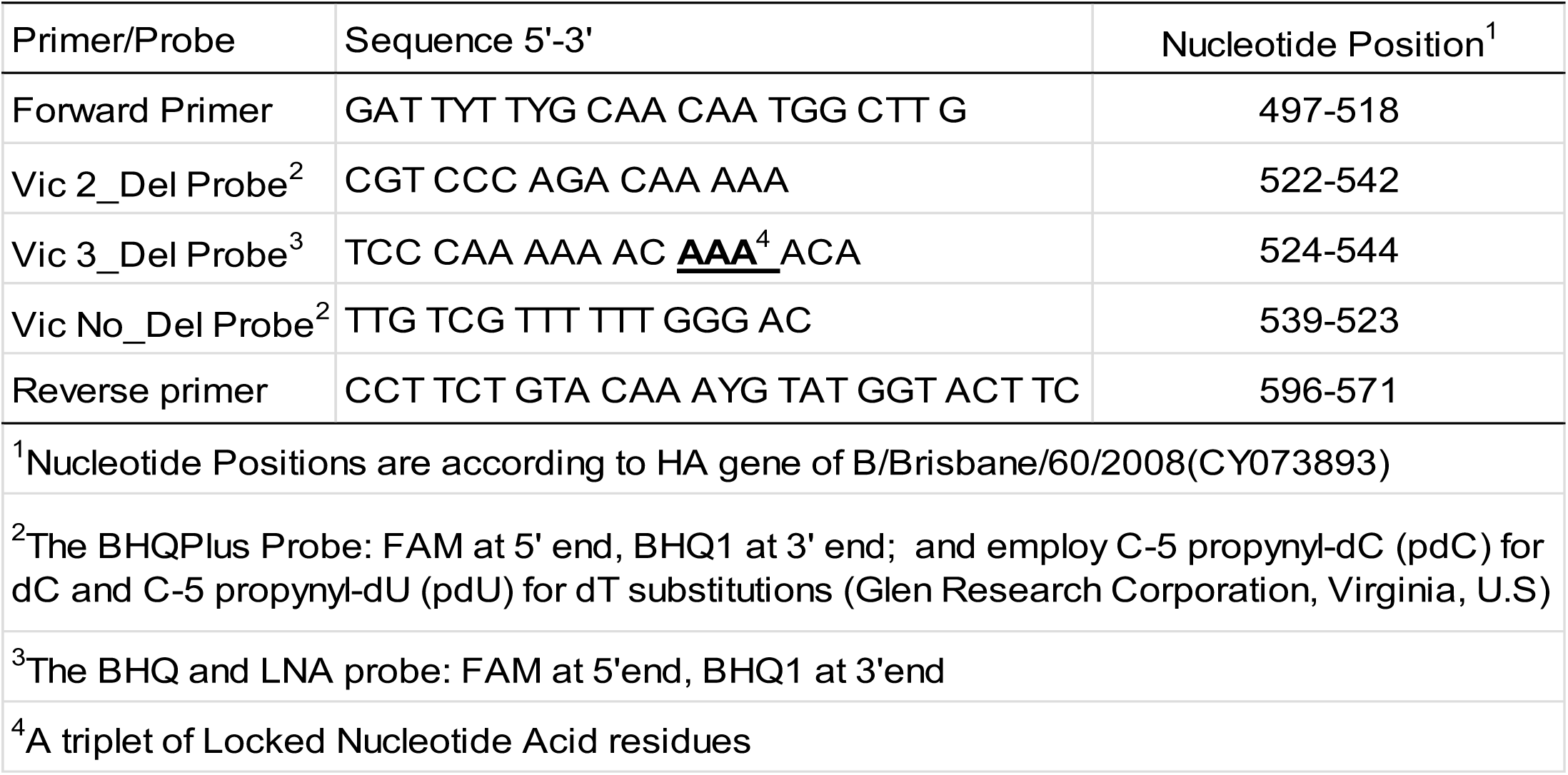
CDC Influenza B/Victoria Lineage Deletion Detection assay primer probe sequences

### rRT-PCR Reaction Conditions

Reaction conditions for rRT-PCR were based upon the FDA-approved CDC Flu rRT-PCR Dx Panel [15, 18]. PCR reaction parameters of the Vic deletion assay were optimized using Invitrogen SuperScript™III Platinum® One-Step quantitative RT-PCR (qRT-PCR) Kits (Single-plex) and TaqPath™ qPCR Multiplex Master Mix (Triplex) kits (Life Technologies) on the Applied Biosystems^TM^ (AB) 7500 Fast Dx Real-Time PCR instrument. All rRT-PCR reactions performed had a total reaction volume of 25µl with primer and probe reaction concentrations at 0.8µM and 0.2µM, respectively. Thermocycling rRT-PCR conditions were as follows: 50°C for 30 min, *Taq* activation at 95°C for 2 min and 45 cycles of 95°C for 15 sec and 55°C for 30 sec. All analytical performance data were collected on an ABI 7500 Fast Dx Real-time PCR instrument. Increases in fluorescent signal were registered during the annealing step of the reaction. All data were analyzed with the sequence detector software (SDS) v1.4.1 (Life Technologies).

### Analytical Sensitivity and Specificity

In order to demonstrate rRT-PCR performance of the VIC deletion assay, five B/Vic lineage viruses and two B/YAM lineage viruses from recent circulating genetic groups, including the WHO recommended influenza B vaccine viruses B/Colorado/6/2017 (V1A.1), B/Brisbane/60/2008 (V1A) and B/Phuket/3073/2013 (B/Yam) were selected for assay evaluation (Figure 1). Analytical sensitivity of the Vic 2_Del and Vic No_Del assays was determined using two V1A.1 and V1A viruses. The Vic 3_Del assay was evaluated by using the V1A.2 virus B/Hong Kong/269/2017 and V1A.3 virus B/Washington/02/2019. The sensitivity of the three assays was evaluated with the commercially available Invitrogen qRT-PCR kit.

The analytical specificity was further assessed using these five B/Vic lineage deletion viruses and two B/Yam lineage viruses, 24 seasonal influenza A and two avian influenza A viruses.

### Assay performance on clinical specimens

To demonstrate the performance of the Vic deletion assay against clinical specimens, 167 clinical specimens including 67 B/Vic, 30 B/Yam and 30 influenza A and B negative, and 30 seasonal influenza A (H1N1pdm-15 and H3N2-15) as determined by the CDC rRT-PCR Flu Panel, were tested with the Vic deletion assay (Table 5).

## RESULTS

### Antigenic analysis

Influenza B viruses from the V1A, V1A.1, V1A.2, V1A.3 and Yam lineages were antigenically characterized in HI tests using post-infection ferret sera raised against representative human Influenza B viruses. HI assays demonstrated that both the V1A.1 V1A.2 and V1A.3 variant influenza B viruses are antigenically distinct from each other and to the no-deletion B/VIC virus (V1A), as the viruses from each of the genetic groups were not well inhibited by antiserum raised against the other genetic groups (reductions in HI titers of 4-32 fold compared to the homologous HI titers) (Table 1).

### Real time RT-PCR assay establishment

The Vic deletion assay is an rRT-PCR assay developed using the ABI 7500 Fast Dx Real-time PCR system, and is used for the qualitative detection and characterization of B/Vic virus RNA in respiratory specimens from patients presenting with influenza-like illness (ILI). The Vic deletion assay employs one pair of conserved primers and three probes specific to each genetic group (Figure 2). The probes target the deletion region of AA position 162-164 (corresponding to nucleotide position 529-537) of the HA gene in influenza B viruses. The AA 162-164 deletion region contains multiple adenines and repeating sequences, thus creating a challenge for designing probes for this region. We found that unmodified RT-PCR probes were unsuccessful in targeting the deletion region due to challenges in the surrounding sequence patterns (data not shown). BHQplus probes contain stabilizing chemistry that allows probe oligonucleotides to be a shorter length which more easily targets regions with challenging sequence patterns [19]. We used BHQplus chemistry to successfully design probes to the V1A and V1A.1 influenza B viruses (Table 2). When designing a probe for detection of the V1A 3DEL viruses, we attempted both a BHQplus design and a Minor Groove Binding (MGB) probe for increased sequence specificity, but neither of these probe modifications were successful (data not shown). Locked Nucleic Acids (LNAs) have been shown to increase stability and sequence mismatch detection [16, 17], and we found that using a triplet of LNAs centered on the mismatch allowed for successful detection of the V1A.2 and V1A.3 viruses.

### Analytical Sensitivity and Specificity

The performance of the Vic deletion assay was evaluated by comparisons to the performance of the universal influenza B assay (InfB) from CDC Flu rRT-PCR Dx Panel. The InfB assay is designed for detection of the nonstructural (NS) gene in all influenza B viruses by targeting highly conserved regions of the NS protein of the influenza B virus.

### Analytical Sensitivity

Analytical performance studies, evaluated by testing ten-fold serial dilutions of RNAs extracted from recently circulating influenza B viruses, demonstrated that the sensitivity of the VIC 2_Del, VIC 3_Del and VIC No_Del assays is comparable to the InfB assay of the CDC Flu rRT-PCR Dx Panel.

The limits of detection (LoD) of the Vic 2_Del, Vic 3_Del and Vic No_Del assays were tested using the three quantified B/Vic viruses B/Maryland/15/2016, B/Hong Kong/269/2017 and B/Florida/103/2016, respectively. The LoD of Vic 2_Del, Vic 3_Del and Vic No_Del assays were 10^1.5^, 10^−0.2^, 10^0.3^ ID_50_/mL, respectively (10^−0.2~1.5^ ID_50_/ml). This correlates to 10^−2.5~ −0.8^ ID_50_ per reaction (5.0 µl/reaction) (Table 3).

**Table 3.** Assay Limit of Detection by using Invitrogen reagents (N=3)

| B/Victoria lineage<br>deletion detection assay | Influenza virus strain tested | Limit of Detection<br>(ID<sub>50</sub>/mL) | Ct Value (mean±SD) |  |
| --- | --- | --- | --- | --- |
|  |  |  | InfB | Deletion assay |
| Vic 2_Del | B/Maryland/15/2016 | 10<sup>1.5a</sup> | 36.14±0.32 | 37.22±0.89 |
| Vic 3_Del | B/Hong Kong/269/2017 | 10<sup>-0.2b</sup> | 33.45±0.38 | 37.31±0.15 |
| Vic No_Del | B/Florida/103/2016 | 10<sup>0.3b</sup> | 35.60±1.16 | 35.35±0.17 |
| <sup>a</sup> Data represent EID<sub>50</sub>/mL; <sup>b</sup> Data represent TCID<sub>50</sub>/mL. |  |  |  |  |

Analytical sensitivity of the Vic deletion assay is shown in Table 4. Three viruses were used to determine LoD of Vic deletion assay and three quantified WHO recommended vaccine component viruses, B/Washington/02/2019 (V1A.3), B/Colorado/6/2017 (V1A.1)) and B/Brisbane/60/2008 (V1A) were evaluated within the lower virus titers, and the results demonstrated Vic 2_Del, Vic 3_Del and Vic No_Del, compared to InfB assay, are sensitive to detect B/Vic deletion variant viruses.

**Table 4.**
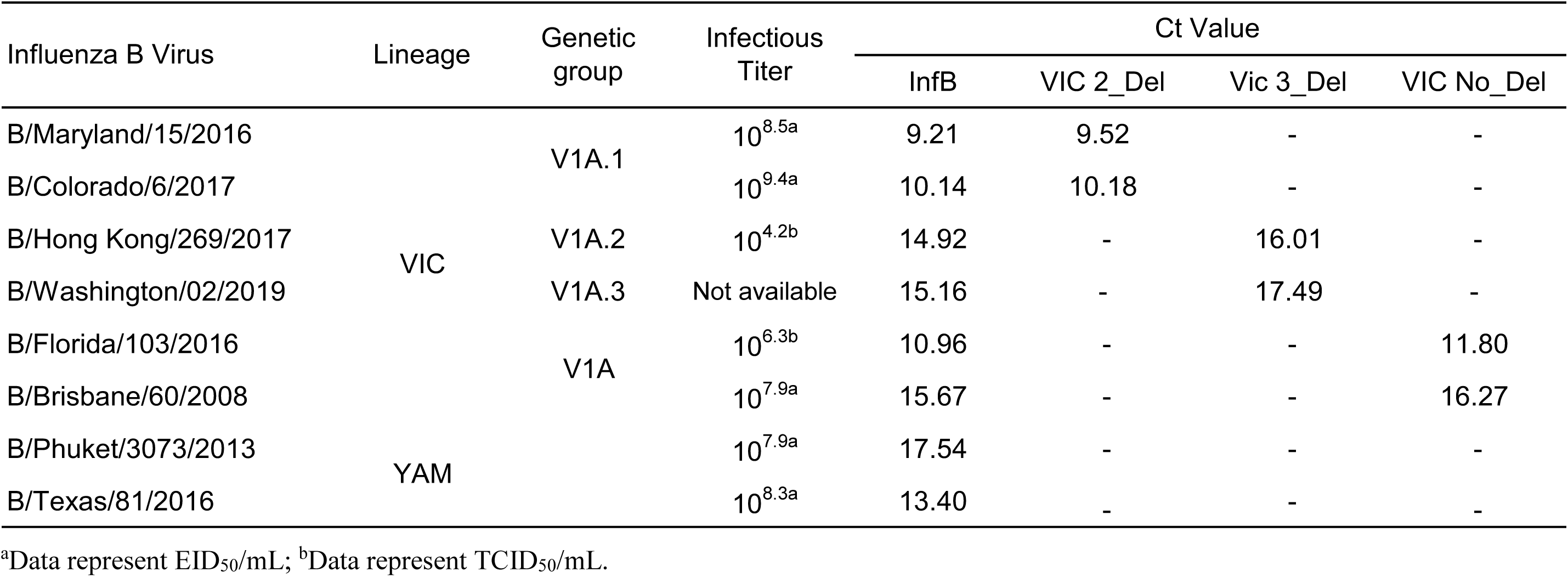
Analytical Specificity (Exclusivity) Testing with Influenza B Victoria and Yamagata lineage Viruses (N=1)

A triplex rRT-PCR assay was developed with one pair of conserved primers with FAM labelled Vic 3_Del probe, Hex labelled Vic 2_Del probe and CAL Fluor Red 610 labelled Vic No_Del probe. This triplex rRT-PCR Vic deletion assay shows the same level of sensitivity and specificity as the single-plex assays in detecting and discriminating the B/Vic deletion variant viruses (Table S1).

### Analytical Specificity (Exclusivity)

Analytical exclusivity was evaluated with high titer influenza B VIC and YAM lineage viruses (Table 4). No cross-reactivity was observed when V1A viruses were tested with the VIC 2_Del and VIC 3_Del assays. Specificity was also demonstrated in V1A.2 and V1A.3 viruses tested with the VIC 2_Del assay and vice versa. Cross reactivity from the VIC No_Del assay was not observed when used to test V1A.1, V1A.2 and V1A.3 viruses. Likewise, the three assays did not react with the B/YAM lineage viruses tested (Table 4).

In order to demonstrate the absence of cross reactivity with influenza A viruses subtypes, exclusively testing was performed by examining twelve each of contemporary seasonal A (H3N2), A/H1N1 pandemic 2009 (A/H1N1pdm09), as well as the benchmark highly pathogenic avian influenza (HPAI) A/H5N1 strain, A/Vietnam/1203/2004, and Asian lineage avian influenza A (H7N9) virus, A/Anhui/01/2013. All influenza A viruses were negative in all three assays (Data not shown).

### Clinical performance of B/Victoria lineage deletion detection assay

To evaluate the clinical performance of the Vic deletion assay, we tested the assay on clinical specimens received at the CDC during influenza surveillance. The Vic deletion assay was successful in classifying the 77 B/Vic clinical specimens into their respective genetic groups (30 V1A-like, 27 V1A.1-like and 20 of V1A.2 and V1A.3-like), and as expected, the rRT-PCR results were confirmed by genetic analysis. All 30 B/Yam, 15 each of A/H1N1pdm09 and A/H3N2, as well as 30 influenza A and B negative samples tested negative for all B/Vic deletion assay targets (Table 5).

**Table 5.**
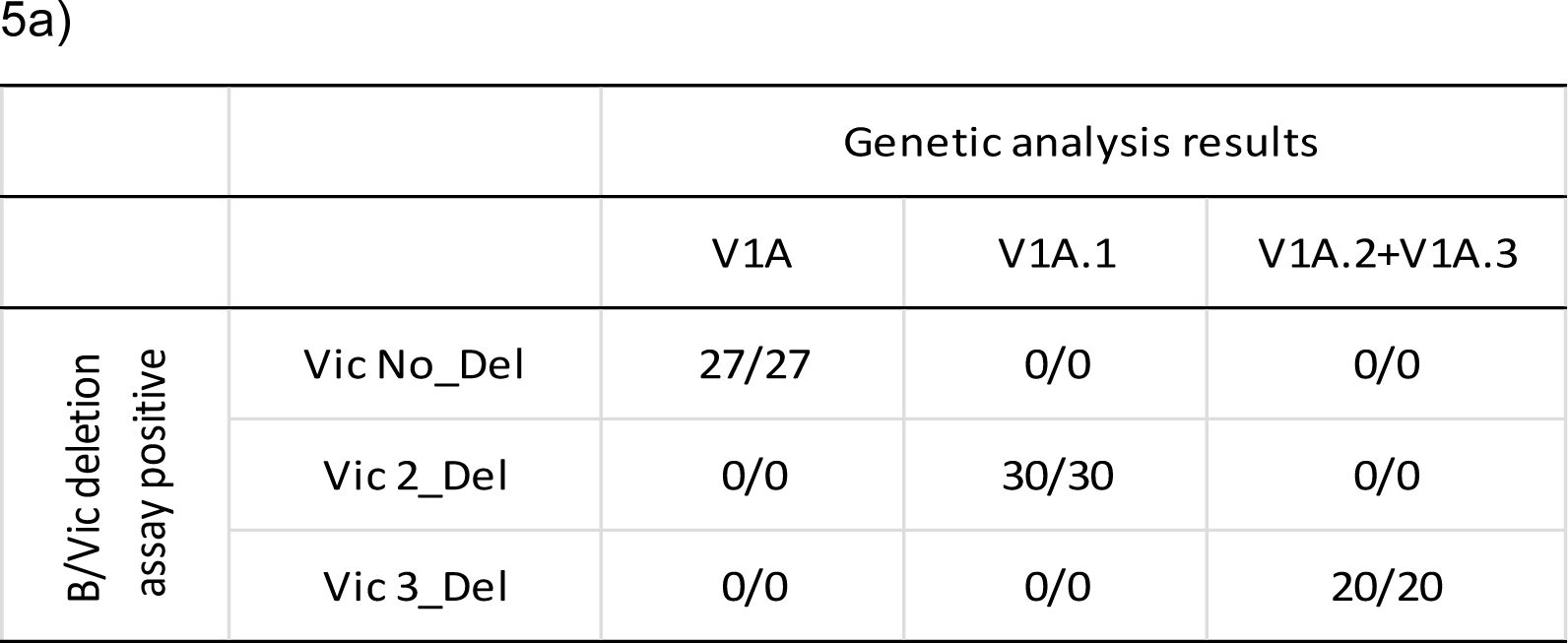
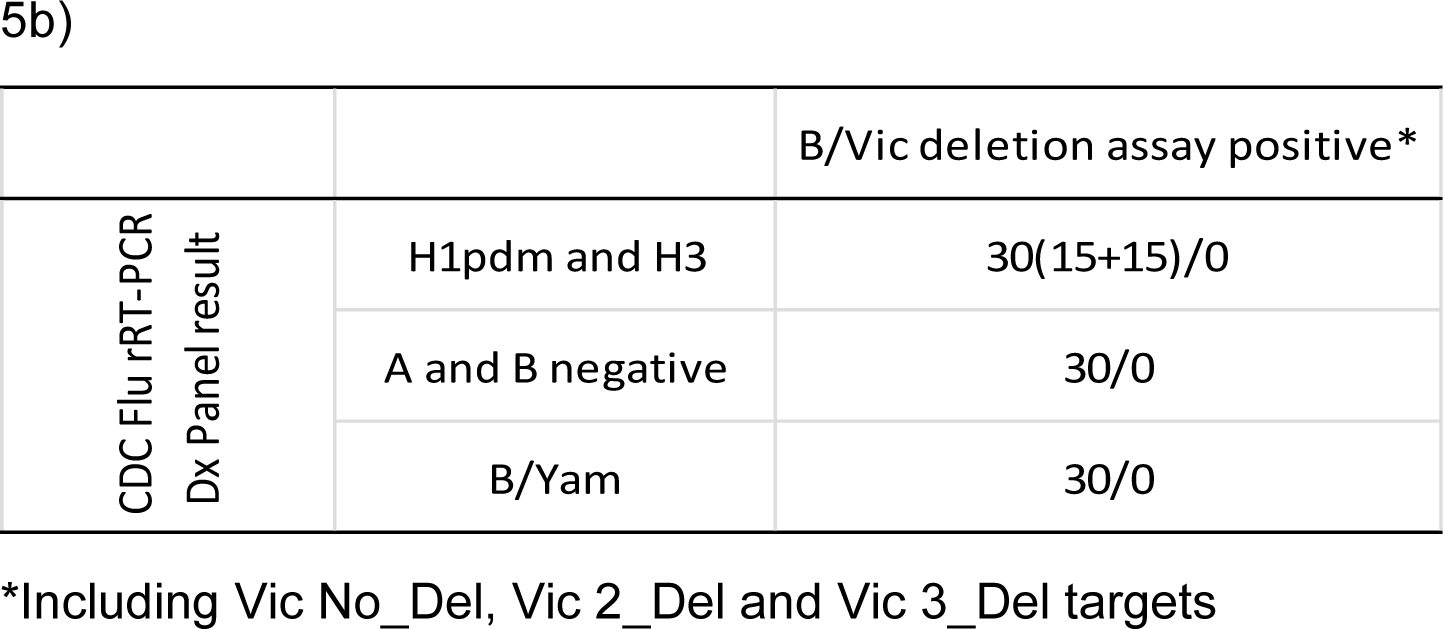
Clinical performance of the B/Victoria Lineage Deletion Detection assay: 5a) Detection of B/Victoria lineage specimens comparing genetic analysis; 5b) Detection of other subtype/lineage influenza specimens

|  |  | Genetic analysis results |  |  |
| --- | --- | --- | --- | --- |
|  |  | V1A | V1A.1 | V1A.2+V1A.3 |
| B/Vic deletion<br>assay positive | Vic No_Del | 27/27 | 0/0 | 0/0 |
|  | Vic 2_Del | 0/0 | 30/30 | 0/0 |
|  | Vic 3_Del | 0/0 | 0/0 | 20/20 |

5b)
|  |  | B/Vic deletion assay positive* |
| --- | --- | --- |
| CDC Flu rRT-PCR<br>Dx Panel result | H1pdm and H3 | 30(15+15)/0 |
|  | A and B negative | 30/0 |
|  | B/Yam | 30/0 |

\*Including Vic No\_Del, Vic 2\_Del and Vic 3\_Del targets

## DISCUSSION

The Vic deletion assay presented here is intended for the qualitative detection of the influenza B/Victoria lineage HA gene deletion variant viruses using rRT-PCR technology. The analytical and clinical performance of the Vic deletion assay in either the single- or tri-plex configuration demonstrates that the assay is highly efficient, sensitive, and comparable to the gold standard CDC rRT-PCR Flu Panel InfB assay[18]. We have further shown that the primer and probes of the VIC deletion assay are specific to V1A, V1A.1 and V1A.2 and V1A.3 viruses, and do not cross react with B/Yam viruses, seasonal influenza A, or avian influenza A viruses including HPAI A/H5N1 and Asian lineage A/H7N9 influenza viruses.

The key genetic distinction between V1A, V1A.1, V1A.2 and V1A.3 viruses is within the same nucleic acid region, presenting a challenge for rRT-PCR probe design. We used two chemical modifications in probe design, LNA and BHQplus chemistry, which allowed for greater stability and specificity. Including LNAs in real-time PCR probes improves detection of mismatches significantly[15, 16]. The Vic 3_Del probe was designed and optimized using a triplet LNA approach, as probes labelled with a triplet of LNA residues centered on the mismatch provide greater discriminatory power than probes with a single LNA modification [16, 17]. The V1A and V1A.1 probes were designed using BHQplus chemistry, allowing for stabilization and enhanced mismatch detection with an overall shorter probe length [19]. We found that unmodified probes were not sufficient for this assay.

During development of the Vic deletion assay, we also evaluated a ZEN_MGB probe that includes an internal ZEN^TM^ quencher located nine nucleotides away from the 5’ FAM reporter dye in addition to a MGB residue quencher at the 3’ end of the probe (Integrated DNA Technologies. Inc. Coralville, IA, U.S). The ZEN_MGB probes performed comparably to the probes presented here in discriminating these influenza B/Vic viral variants [20]. Thus, both BHQplus and ZEN_MGB fluorescent hydrolysis probe quencher chemistries can be used to synthesize probes for the Vic deletion assay.

Sequence alignments using HA gene sequences from 2010-2019 demonstrated the conserved primers to be stable with no conserved genetic changes observed (Data not shown). Although the areas chosen for the conserved amplification primers and the deletion type-specific probes are currently stable, genetic changes due to rapidly virus evolution and the variable nature of RNA viruses may require periodic updates of the B Vic deletion assay primer and probe sequences.

The number and percentage of circulating V1A.1 V1.2 and V1A.3 viruses has increased significantly since initial identification [21], making it critical to have a simplified and sensitive assay capable of detecting these genetic variants during influenza surveillance. The Vic deletion rRT-PCR assay is the most sensitive method currently available that can distinguish these influenza B genetic subgroups. Recently, a pyrosequencing method was published that can also be used to distinguish these influenza B genetic variants[10]. However, the pyrosequencing method requires costly instruments that are not common to diagnostic laboratories, and rRT-PCR is more sensitive than sequencing-based methods [22, 23]. A conventional RT-PCR assay has also been recently developed to distinguish these influenza B genetic subgroups, but conventional RT-PCR is less sensitive and more time consuming than a real-time RT-PCR assay[11]. Given its sensitivity and specificity, the rapidity of results, the fact that the Vic deletion assay can be run on equipment that is readily available in most diagnostic labs, and the fact that the assay can be multiplexed for testing efficiency and cost-savings, this is the most optimal assay for influenza surveillance currently available. Combining the CDC Vic deletion assay with the CDC Flu rRT-PCR Dx Panel will allow for rapid identification of all five of the distinct influenza B viral genetic groups, including B/Yam, B/Vic V1A, V1A.1, V1A.2 and V1A.3 viruses. This will allow for understanding the health burden of each of these antigenically and genetically distinct viruses in different global regions, which in turn, will allow for the most relevant vaccine virus recommendations moving forward into future influenza seasons.

## Supplemental materials

**Table S1.** Analytical sensitivity of the B/Victoria Lineage Deletion Detection assay single- and multi-plex assays: Vic 2_Del assay against B/Vic V1A.1 viruses (S1a), Vic 3_Del assay against V1A.2 viruses (S1b) and Vic No_Del assay against V1A-the other B/Vic viruses (S1c) (N=3)

| Influenza B V1A.1 viruses<br>(EID<sub>50</sub>/mL) | Ct Value (Mean±SD) |  |  |
| --- | --- | --- | --- |
|  | InfB | Vic 2_Del |  |
|  |  | Single (FAM) | Triplex(Hex) |
| B/Maryland/15/2016 |  |  |  |
| 10<sup>4.5</sup> | 26.17±0.32 | 24.53±0.36 | 26.13±0.04 |
| 10<sup>3.5</sup> | 29.60±0.91 | 28.70±0.14 | 29.86±0.12 |
| 10<sup>2.5</sup> | 35.19±1.06 | 32.60±0.26 | 34.20±0.26 |
| B/Colorado/06/2017 |  |  |  |
| 10<sup>4.4</sup> | 29.48±0.43 | 27.95±0.23 | 29.26±0.21 |
| 10<sup>3.4</sup> | 33.54±0.28 | 32.38±0.10 | 33.57±0.09 |
| 10<sup>2.4</sup> | 37.13±0.97 | 35.84±1.31 | 36.28±0.03 |

S1b
| Influenza B V1A.2 virus<br>(TCID<sub>50</sub>/mL) | Ct Value (Mean±SD) |  |  |
| --- | --- | --- | --- |
|  | InfB | Vic 3_Del |  |
|  |  | Single (FAM) | Triplex(FAM) |
| B/HongKong/269/2017 |  |  |  |
| 10<sup>2.2</sup> | 22.93±0.51 | 26.37±1.29 | 26.48±0.18 |
| 10<sup>1.2</sup> | 27.64±0.79 | 28.85±0.65 | 30.12±0.15 |
| 10<sup>0.2</sup> | 31.70±0.77 | 34.41±1.75 | 33.63±0.35 |

S1c
| Influenza B V1A viruses<br>(ID<sub>50</sub>/mL) | Ct Value (Mean±SD) |  |  |
| --- | --- | --- | --- |
|  | InfB | Vic No_Del |  |
|  |  | Single(FAM) | Triplex(FAM) |
| B/Florida/103/2016 |  |  |  |
| 10<sup>3.3a</sup> | 26.23±0.59 | 24.06±0.09 | 25.10±0.13 |
| 10<sup>2.3</sup> | 30.53±0.56 | 28.38±0.09 | 29.94±0.60 |
| 10<sup>1.3</sup> | 34.77±0.98 | 31.80±0.45 | 33.47±0.36 |
| B/Brisbane/60/2008 |  |  |  |
| 10<sup>4.9b</sup> | 29.47±0.55 | 26.65±0.28 | 28.38±0.12 |
| 10<sup>3.9</sup> | 34.42±0.97 | 31.14±0.06 | 32.70±0.37 |
| 10<sup>2.9</sup> | 36.79±0.19 | 34.54±0.27 | 37.29±0.26 |

<sup>a</sup> Data represent TCID<sub>50</sub>/mL; <sup>b</sup>Data represent EID<sub>50</sub>/mL.

**Figure S1.**
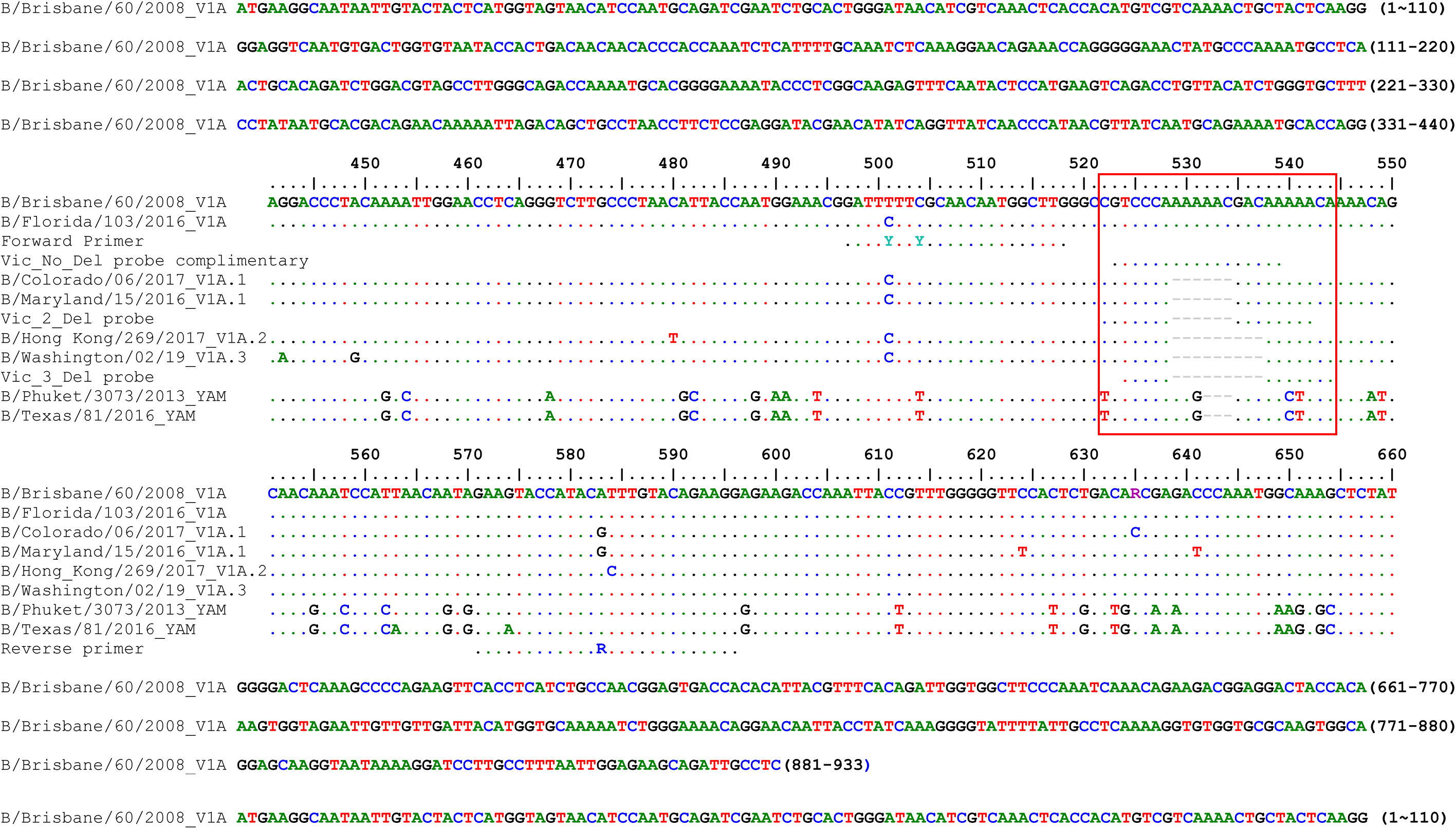
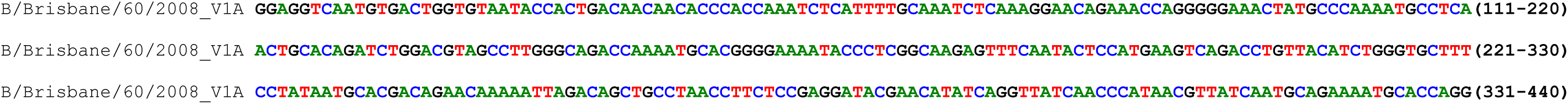
Primer Probe sequences aligned with reference viruses. Conserved primers and genetic group specific probes, targeting on deletion region of amino acid position 162-164 (corresponding to nucleotide position 529-537; red box) of HA gene (HA1 region), were designed to specifically detect and differentiate influenza B Victoria lineage deletion variant viruses

